# Bacteriophage-derived dsRNA exerts anti-SARS-CoV-2 activity *in vitro* and in Golden Syrian hamsters *in vivo*

**DOI:** 10.1101/2022.01.22.477073

**Authors:** Vaivode Kristine, Verhovcova Irina, Skrastina Dace, Petrovska Ramona, Kreismane Madara, Lapse Daira, Rubene Diana, Kalnina Zane, Pjanova Dace

## Abstract

**Purpose:** Bacteriophage-derived dsRNA, also known as Larifan, is nationally well-known broad-spectrum antiviral medication. The goal of this study was to ascertain the antiviral activity of Larifan against the novel SARS-CoV-2.

**Methods:** The antiviral activity of Larifan against SARS-CoV-2 *in vitro* was measured in human lung adenocarcinoma (Calu3) and primary human small airway epithelial cells (HSAEC) using cytopathic effect assay, viral RNA copy number detection by digital droplet PCR (ddPCR) and infectious virus titration in cells supernatants in Vero E6 cells by end-point titration method. The antiviral effect of Larifan *in vivo* was detected in SARS-CoV-2 infection model in Golden Syrian hamsters. Larifan (5 mg/kg) was administered either subcutaneously or intranasally twice before and after virus infection with a 24-hour interval between doses. The viral RNA copies and infectious virus titre were detected in animal lungs at day three and five post-infection using ddPCR and end-point titration in Vero E6 cells, respectively. Histopathology of lungs was analysed as well.

**Results:** Larifan inhibited SARS-CoV-2 replication in Calu3 cells both after the drug addition pre- and post-infection with a substantial drop in the supernatant viral RNA copy numbers from eight (p = 0.0013) to twenty (p = 0.0042) times, respectively. Similarly, infectious virus titre in Vero E6 cells dropped by 3.6log_10_ TCID_50_ and 2.8log_10_ TCID_50_ after the drug addition pre- and post-infection, respectively. In HSAEC, Larifan inhibited SARS-CoV-2 replication at the similar level. Larifan also markedly reduced virus numbers in the lungs of infected hamsters (p = 0.0032) both at day three and five post-infection with a more pronounced effect after intranasal administration reaching a drop by 2.7log_10_ at day three and 2.0log_10_ at day five. The administration of Larifan also reduced the amount of infections virus titer in lungs (p = 0.0039) by 4.3log_10_ TCID_50_ and 2.8log_10_ TCID_50_ at day three and five post-infection, respectively. Improvements in the infection-induced pathological lesion severity in the lungs of animals treated with Larifan were also demonstrated by histological analyses.

**Conclusions:** The inhibition of SARS-CoV-2 replication *in vitro* and the reduction of the viral load in the lungs of infected hamsters treated with Larifan alongside the improved lung histopathology, suggests a potential use of Larifan in controlling the COVID-19 disease in humans.

## Introduction

In December 2019, a series of pneumonia cases of unknown cause emerged in Wuhan, China, with clinical presentations greatly resembling viral pneumonia. This disease was caused by a novel form of coronavirus named ‘severe acute respiratory syndrome coronavirus 2’ (SARS-CoV-2), and the disease was accordingly named the coronavirus disease 2019 (COVID-19). On March 11, 2020, World Health Organisation announced this emerging disease as a fast-progressing pandemic. COVID-19 is a complex disorder, and its manifestations can range from mild upper respiratory illness to severe bilateral pneumonia, acute respiratory distress syndrome, disseminated thrombosis, multi-organ failure and death (1,2).

Since the emergence of this disease, numerous anti-COVID-19 vaccines have been tested in clinical trials worldwide. Several have been approved and are currently successfully used to immunize people against this novel virus. Nonetheless, antiviral drugs slowing down the replication of SARS-CoV-2 could be beneficial and fulfil an essential role in treating COVID-19 patients, *e*.*g*. preventing or alleviating disease symptoms. In addition, antiviral drugs could be used as a preventive measure to protect high-risk groups, especially if they could be administered in a non-invasive manner.

Here, we present a nationally well-known antiviral drug in the form of double-stranded RNA (dsRNA) as a potential anti-COVID-19 medication. Bacteriophage-derived dsRNA, also known as Larifan (Larifan Ltd., Riga, Latvia), is a heterogeneous population of dsRNAs. It has been obtained biotechnologically from *E. coli* cells infected with f2sus11 amber mutant bacteriophage, and comprises of dsRNA molecules (acidum ribonucleinicum duplicatum) with an average length of 700 base pairs. Larifan has been developed as a poly-functional and wide-spectrum antiviral drug shown to be a potent inducer of endogenous type I interferons (IFNs). At the time of its invention, the mechanism of the Larifan’s action was studied by Sokolova et al. in both experimental systems (3) and clinical trials on volunteers (4). The lateral results proved Larifan’s ability to induce and activate enzymes of the IFN system, which are involved in the translation blockage in virus-infected cells (5). In the present study, we evaluated the antiviral efficiency of Larifan against SARS-CoV-2 *in vitro* in human lung adenocarcinoma cell line (Calu3), primary human small airway epithelial cells (HSAEC) and *in vivo* in an infection model of Golden Syrian hamsters. Promisingly, Larifan exserts an antiviral effect against SARS-CoV-2 *in vitro* and *in vivo*.

## 2. Materials and methods

### SARS-CoV-2

The SARS-CoV-2 strain used in this study, SARS-CoV-2 hCoV-19/Sweden/20-53846/2020 (lineage B 1.1.7, UK), was obtained from European Virus Archive and propagated in Vero E6 cells; passage four virus was used for the studies described here. The titre of the virus stock was determined by 50% tissue culture infective dose (TCID_50_) according to the cytopathic effect using the Reed-Muench method. All the infection experiments were performed in a biosafety level-3 (BSL3) laboratory.

### Cell lines

Vero E6 cells (African green monkey kidney, ATCC CRL-1586) and Calu3 (human lung adenocarcinoma cells, ATCC HTB-55) were cultured in Dulbecco’s Modified Eagle Medium (DMEM) supplemented with 10% fetal bovine serum (FBS), 1% L-glutamine and 1% bicarbonate (all from Gibco). HSAEC (primary human small airway epithelial cells, ATCC PCS-301-010) were grown in Airway Epithelial Cell Basal Medium (ATCC PCS-300-030) supplemented with Bronchial Epithelial Cell Growth Kit (ATCC PCS-300-040). Virus end-point titrations in Vero E6 cells were performed with medium containing 2% FBS instead of 10%. All cells were incubated in a humidified atmosphere at 37°C with 5% CO_2_.

### Cytotoxicity assay

The cytotoxic effects of Larifan on Calu3 and HSAEC cells were evaluated by the Cell Counting Kit-8 (CCK8 – Dojindo Laboratories). Monolayers of Calu3 and HSAEC cells in 96-well plates were incubated with indicated concentrations of Larifan. After 72 h, CCK8 solution was added, and cells were incubated for additional 4 h at 37°C with 5% CO_2_. The absorbance was measured at 450 nm. Half-maximal cytotoxic concentration (CC_50_) was calculated according to the best fit point-to-point line’s interpolated value.

### Infection of cells with SARS-CoV-2 and treatment with Larifan

Calu3 and HSAEC cells were grown to a monolayer in 6-well plates. Cells were infected with SARS-CoV-2 at a multiplicity of infection of 0.3. Samples for virus assessment, including cell culture supernatants were collected every 24 h. Cells were treated with varying concentrations of Larifan. Cells were pre-treated with Larifan for 4 h, and the virus was then added to allow attachment for 1 h. Afterwards, cells were cultured with a drug-containing medium until the end of the experiment (“full-time treatment”). For pre-infection treatment, Larifan was added to the cells for 4 h before viral attachment. Then cells were cultured in fresh culture medium without the drug. For post-infection treatment, Larifan was added immediately after virus infection and maintained in the medium until the end of the experiment. Following 72 h of incubation, samples were collected for virus presence quantification by end-point titration in Vero E6 cells and ddPCR for infectious virus and virus copy number detection, respectively.

### SARS-CoV-2 infection model in hamsters

We have used the hamster infection model of SARS-CoV-2 as described previously (6). The experimental procedures in animals were approved by the National animal welfare and ethics committee (permit no. 124/2021) and were performed in compliance with the Directive 2010/63/EU as adopted in the national legislation.

In total, 24 naïve 9-10 weeks old specific pathogen free (SPF) male Golden Syrian hamsters (strain HsdHan^®^:Aura) were purchased from Envigo (US) and after introduction they were single-housed in individually ventilated cages GR900 (Tecniplast, Italy), HEPA-ventilated by SmartFlow air handling unit (Tecniplast, Italy) at 75 air changes per hour in a negative pressure mode. Access to autoclaved water acidified to pH 2.5-3.0 with hydrogen chloride and standard rodent diet (4RF21 (A), Mucedola) was provided *ad libitum*. Aspen wooden bedding and nesting material (Tapvei, Estonia) together with rat cardboard houses (Velaz, Czech Republic) and aspen gnawing bricks (Tapvei, Estonia) were provided in all cages. Animals were housed in SPF facility BSL3 unit under controlled temperature (24±1°C) and relative humidity of 40-60%. All animals were subjected to at least 7-day acclimatization period with 7:00 am–7:00 pm visible light cycle. An individual animal served as an experimental unit in these experiments.

For the Covid-19 modelling, the isoflurane-anesthetised animals were intranasally infected with SARS-CoV-2. Briefly, animals were induced with 5% isoflurane and maintained under 2% isoflurane while inoculated intranasally with 100 μl containing 2×10^4^ TCID_50_ of the virus (50 μl per nostril).

Animals received the drug treatment either subcutaneously (s.c.) or intranasally (i.n.) using Larifan dosage 5 mg/kg in phosphate-buffered saline. Hamsters were pre-treated with Larifan twice before virus infection (the second drug administration four hours before infection) and twice after virus infection with an interval of 24 h between each drug administration. Each experimental group consisted of eight hamsters. Group size was estimated using resource equation approach (7). Hamsters were daily monitored for clinical signs, appearance, behaviour, and weighted. At days three and five four hamsters from each group were humanely euthanized using 5% isoflurane anaesthesia. Lungs were collected for histological observations and virus quantification by ddPCR, and end-point titration.

### RNA isolation and viral RNA quantification by digital-droplet PCR (ddPCR) analysis

Viral RNA from cell culture supernatants was isolated as the manufacturer’s instructions using QIAamp MinElute Virus kit (Qiagen). Cells were suspended in TRI Reagent (Sigma Aldrich), and total RNA was extracted according to manufacturer’s protocol. Hamster lungs were collected and homogenized using bead disruption with Lysing Matrix E from MP Biomedicals (US) either in TRI Reagent or DMEM for virus copy number and infectious virus detection, respectively, and centrifuged to pellet cell debris. Viral copy numbers were assessed via digital-droplet PCR (ddPCR) using SARS-CoV-2 ddPCR kit (#12013743) from Bio-Rad with N1 and N2 primers and probes targeting the nucleocapsid genes. Droplets were generated using Bio-Rad QX200 Droplet Generator and analyzed with QX200 Droplet Reader (Bio-Rad). Absolute quantifications of virus RNA copies were estimated by modelling a Poisson distribution using QuantaSoft™ analysis software version 1.7 (Bio-Rad). Mean signal from N1 and N2 was extrapolated and expressed as viral RNA copies per microliter of supernatant, number of cells or milligrams of tissue.

### End-point virus titration

Cell culture supernatants and lung homogenates were put directly onto Vero E6 cells. End-point titrations were performed on confluent Vero E6 cells in 96-well plates. Viral titres were calculated by the Reed-Muench method and expressed as TCID_50_ per microliter of supernatant, number of cells or milligrams of tissue.

### Histology

The lungs were fixed in 4% formaldehyde immediately *post-mortem* and embedded in paraffin. Tissue sections (7 μm) were stained with hematoxylin and eosin stains and analyzed blindly for lung damage by a certified pathologist.

### Statistical analysis

GraphPad Prism (GraphPad Software, Inc.) was used to perform statistical analysis. Statistical significance was determined using an Unpaired t-test for the *in vitro* experiments and Two-way ANOVA when comparing the *in vivo* groups. P values of ≤ were considered significant.

## 3. Results

### Replication of SARS-CoV-2 in Calu3 and HSAEC cells

Cytolysis of cells was evident only in Calu3 cells (Fig. 1A). Viral RNA copy numbers in the supernatant of Calu3 cells were abundant and peaked on days three and four post-infection (Fig. 1C). In HSAEC cells, SARS-CoV-2 infection did not elicit the typical cytopathic effect (CPE) (Fig. 1B) that is characteristic for many respiratory viruses; however, cells appeared morphologically to “inflate”, and virus release within supernatant could be measured with the increase of viral RNA copy numbers even until day six (Fig. 1D). This suggests a different replication mode of SARS-CoV-2 in epithelial cells that does not produce apparent damage to the cells but allows continuous virus release.

**Figure 1.**
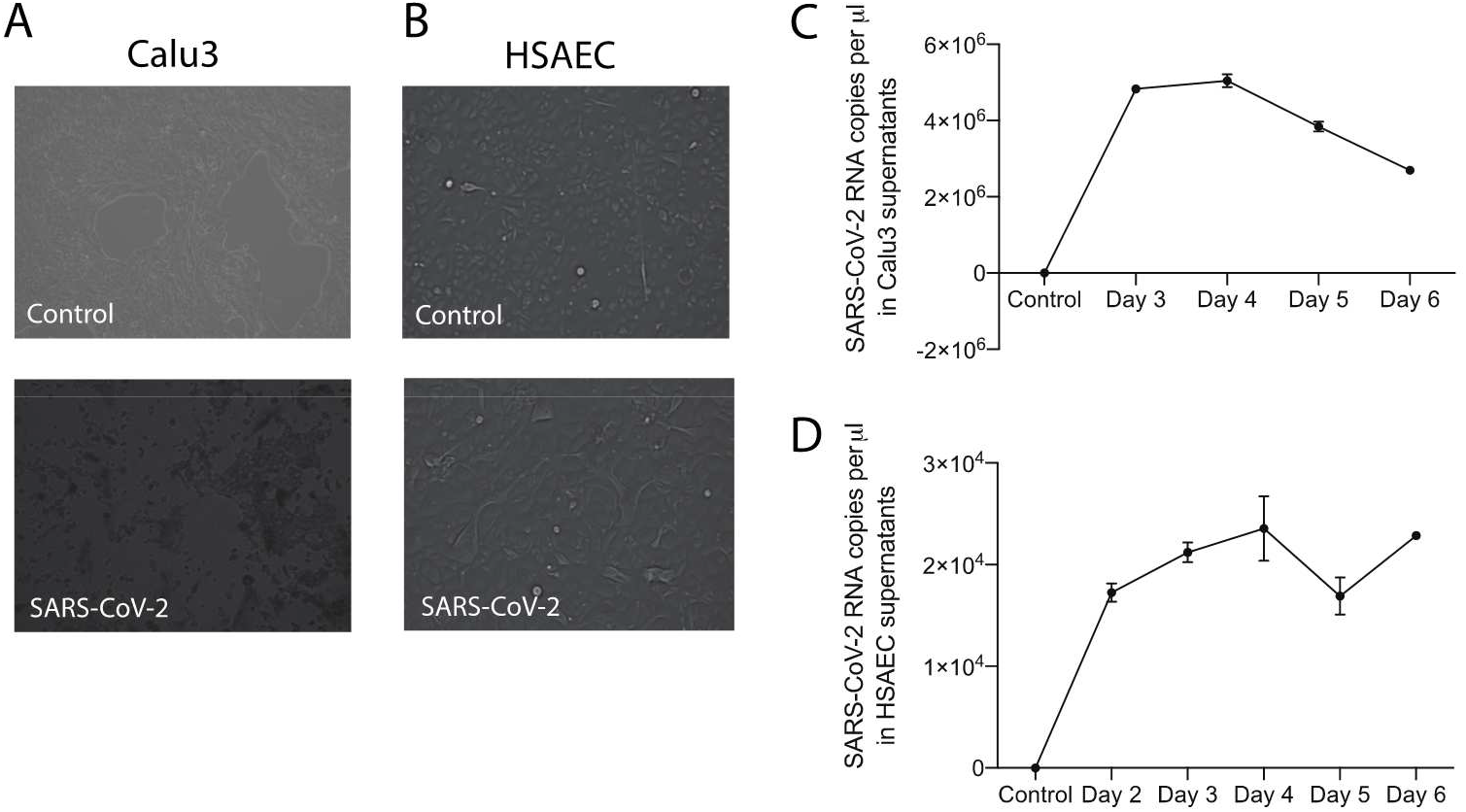
SARS-CoV-2 replication *in vitro* in Calu 3 cells and HSAEC. **(A)** CPE observed in infected Calu3 cells at day three post-infection. **(B)** Morphology of HSAEC at the day three post-infection; no CPE observed. **(C, D)** Viral RNA copy number measurements performed in duplicate in Calu3 cell and HSAEC supernatants during experiment using ddPCR. Control represents uninfected cells.

### Larifan exserts an antiviral activity against SARS-CoV-2 *in vitro*

Larifan showed unapparent cytotoxicity in Calu3 and HSAEC cell lines at all the concentrations used in this study (Fig. 2A) and inhibited the replication of SARS-CoV-2. Viral RNA copy numbers in the supernatant of Calu3 cells dropped from 8.3×10^6^ to an average of 1.1×10^6^ pre-infection (p = 0.0013) and from 13.7×10^6^ to an average of 6.9×10^5^ post-infection (p = 0.0042) after the cells were treated with Larifan (Fig. 2B). Titre of the infectious 7.3log_10_ TCID_50_ virus also dropped significantly in Calu3 cell supernatant before infection to an average of 3.7log_10_ TCID_50_ (p = 0.0094) and to 4.5log_10_ TCID_50_ after infection (p = 0.0186) (Fig. 2C). A comparable reduction in viral RNA copy numbers was observed when Larifan was present throughout (“full-time treatment”) in HSAEC (Fig. 2D).

**Figure 2.**
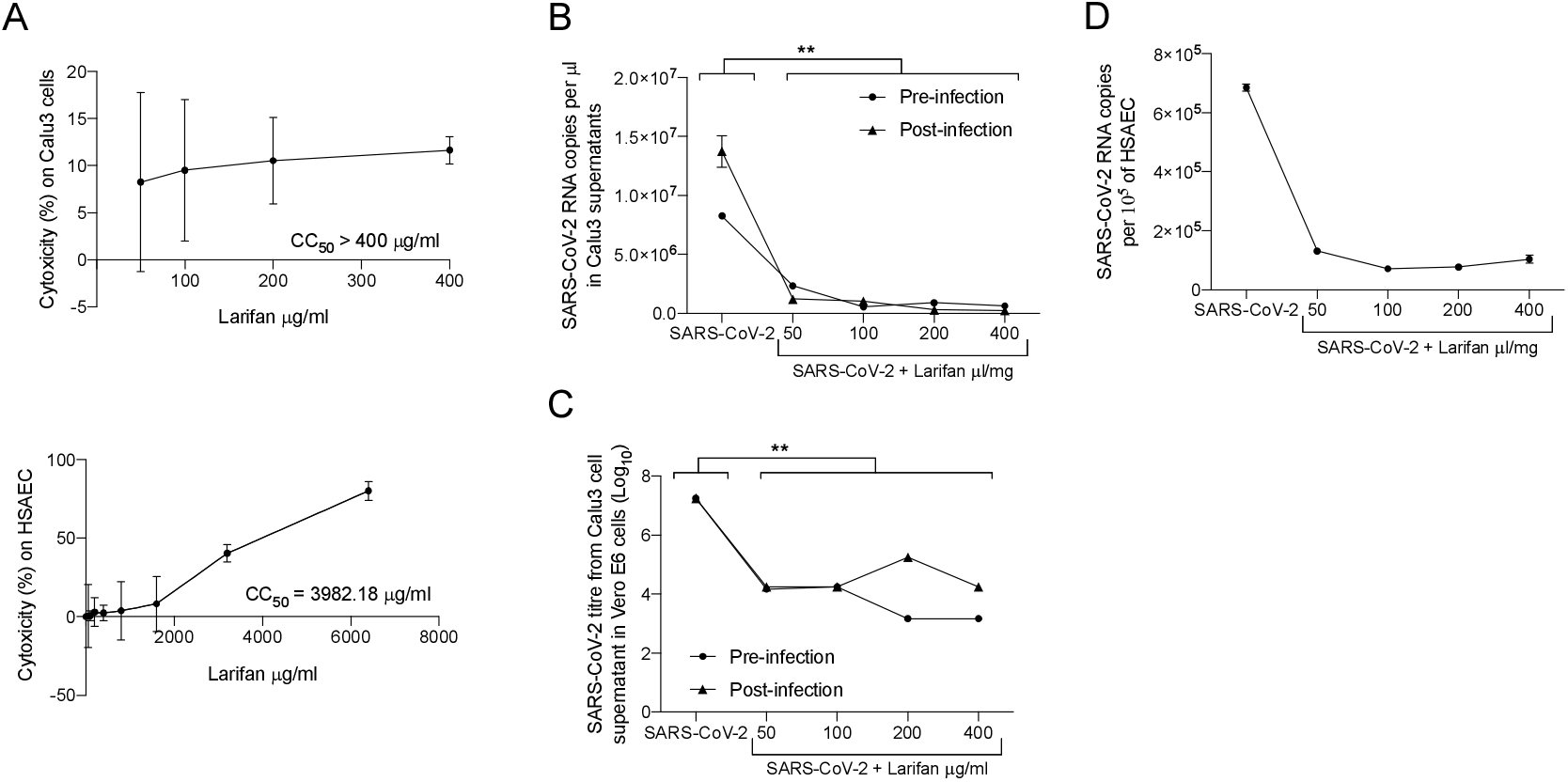
The antiviral activity of Larifan against SARS-CoV2 *in vitro*. **(A)** No significant cytotoxicity of Larifan observed in Calu3 cells or HSAEC done in triplicate. **(B)** Viral RNA copy number drop in Calu3 cell supernatants at day three post-infection when Larifan was added before or after infection done in duplicate. **(C)** Infectious SARS-CoV-2 titre drop in Calu3 cell supernatants in paired experiments in Vero E6 cells after treatment with Larifan at day three post-infection when Larifan was added before or after infection. **(D)** Viral RNA copy number drop in a single measurement in HSAEC when Larifan was present throughout.

### Administration of Larifan decreases SARS-CoV-2 virus numbers *in vivo*

The experimental strategy of the study is depicted in Figure 3A. Hamsters treated with Larifan either s.c. or i.n. presented a decrease in both viral RNA copies and infectious virus titre per mg of lung tissue compared to untreated hamsters. Such an effect was observed at day three and five post-infection, with more pronounced differences at day three. Intranasal Larifan administration gives a reduction of 2.7log_10_ and 2.0log_10_ RNA copies per mg of lung tissue at the days three and five post-infection, respectively, which significantly differs from the viral numbers measured without the presence of Larifan (p = 0.0032) (Fig. 3B). Infectious virus titres in the lung also dropped by 4.3log_10_ TCID_50_ at day three, and by 2.8log_10_ TCID_50_ at the day five post-infection, which also significantly differed from the untreated group (p = 0.0039) (Fig. 3C). Similarly, Larifan administration s.c. also resulted in the detection of fewer virus RNA copies and smaller virus titres in the lung. Although, these changes were less pronounced with the drop of 1.7log_10_ RNA copies per mg of lung tissue at day three post-infection and no changes were observed at day five (Fig. 3B). The drop of infectious virus titres was less pronounced in the case of s.c. administration and was 2.3log_10_ TCID_50_ at day three and 1.5log_10_ TCID_50_ at day five (Fig.3C). A more pronounced effect was measured when Larifan was administrated i.n. reaching statistical significance between administration modes, particularly in the number of RNA copies measured (p = 0.027) (Fig. 3B).

**Figure 3.**
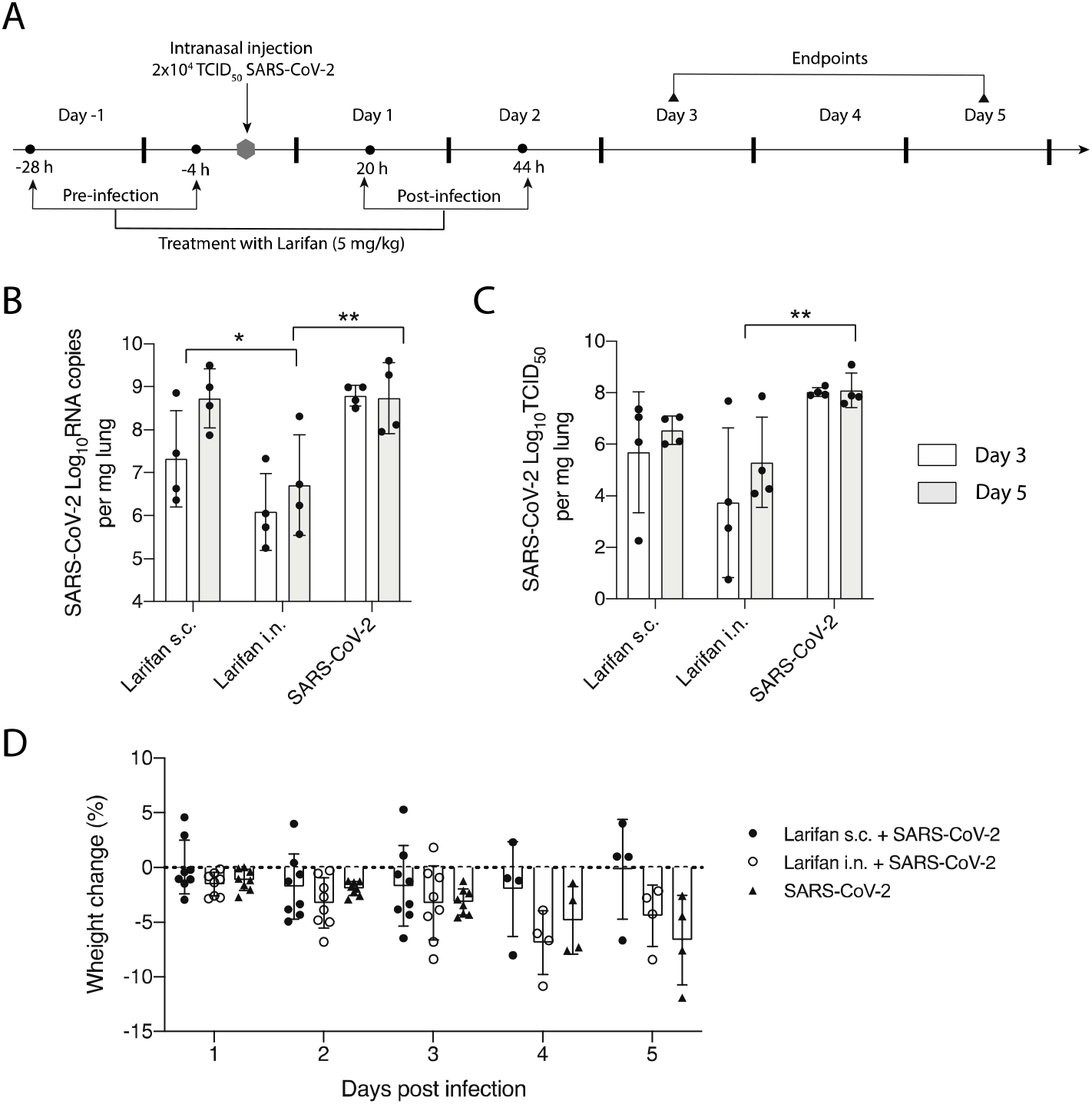
*In vivo* testing of Larifan’s effect in the SARS-CoV-2 infection model of Golden Syrian hamsters. **(A)** Experimental strategy. **(B)** Viral RNA copy number in the lungs of uninfected and SARS-CoV-2 infected hamsters at day three and five after s.c. or i.n. Larifan administration. **(C)** Infectious virus titre in the lungs of uninfected and SARS-CoV-2 infected hamsters at day three and five after s.c. or i.n. Larifan administration. **(D)** Weight changes of uninfected and SARS-CoV-2 infected hamsters during the study.

Intranasal administration of Larifan caused slightly higher weight loss in treated animals during the first days after infection than untreated animals. However, on day five, the opposite effect was observed, and Larifan treated hamsters had lost less weight when compared to untreated animals (Fig. 3D). However, these changes were not statistically significant and might relate to Larifan’s systemic effect on the living organism as it mimics viral infection. Weight gain on day five might indicate faster recovery. Subcutaneous administration of Larifan caused minor weight loss and was comparable with that of untreated hamsters or even less. On day five, the weight loss of treated animals was noticeably smaller than that of untreated hamsters (Fig. 3D).

### Administration of Larifan improves histological lung pathology

In Larifan treated hamsters, improvements in the infection-induced pathological lesion severity in the lungs was also observed (Fig. 4). Infected hamsters’ lungs show several pathological changes: i) alveolar septum fibrosis, ii) alveolar lumens were filled with inflammatory cells and erythrocytes, and iii) peribronchial inflammation could be observed. Noteworthy, in the lungs of Larifan treated hamsters, these changes were less pronounced. On day three, in the lungs of treated hamsters’ pathological changes were observed rarely and the lungs overall had a more healthy appearance. On day five, the inflammation was mainly focal and thus less pronounced.

**Figure 4.**
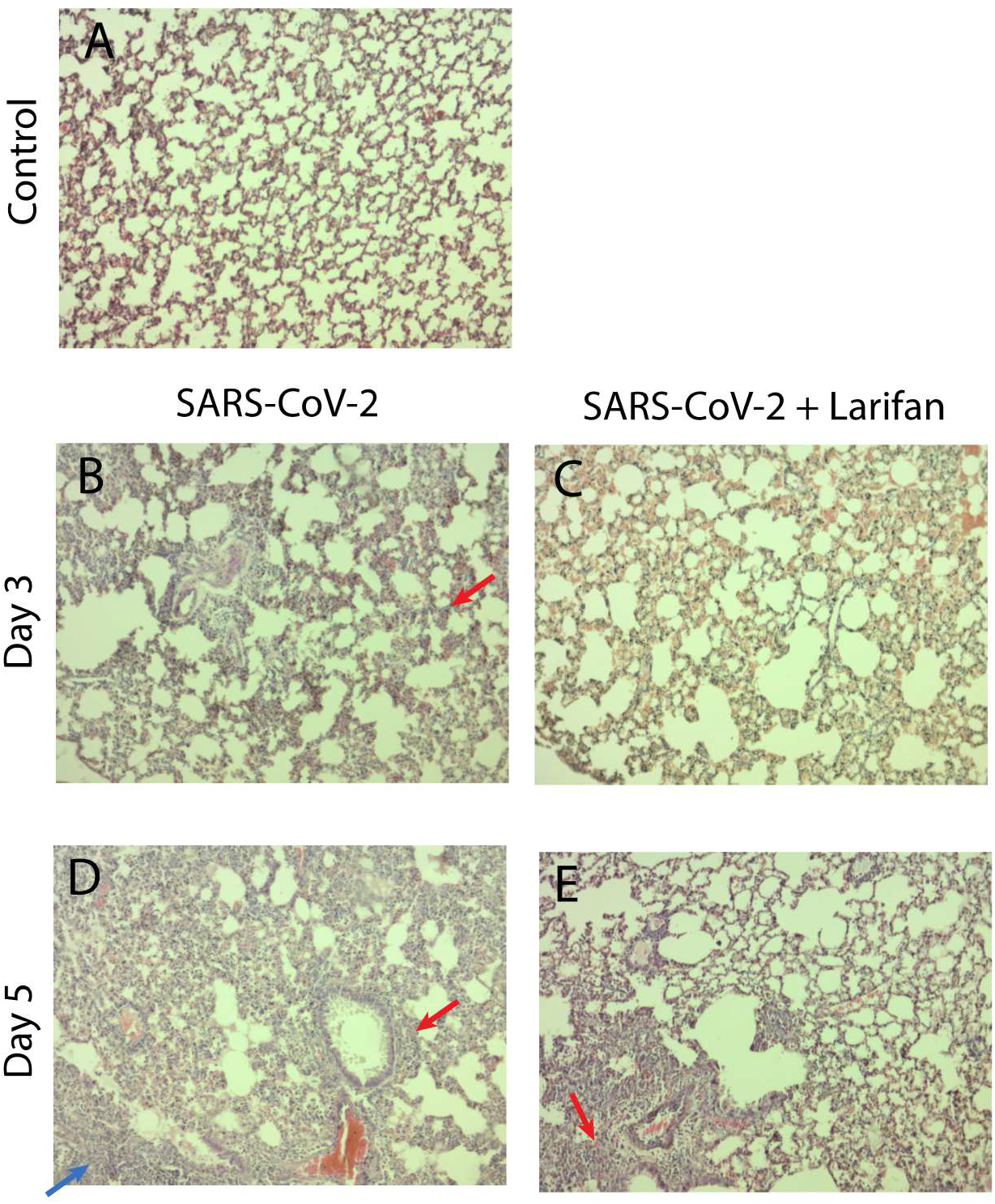
Representative hematoxylin and eosin stained images of lungs of SARS-CoV-2 infected hamsters after s.c. Larifan administration. **(A)** Normal lungs from uninfected control hamster. **(B)** Alveolar septum fibrosis (red arrow) in lungs of untreated hamster three days post-infection. **(C)** Lungs of Larifan treated hamster three days post-infection. **(D)** Lungs of untreated hamsters five days post-infection – alveolar lumens filled with inflammatory cells and erythrocytes (blue arrow), peribronchial inflammation (red arrow). **(E)** Lungs of Larifan treated hamster five days post-infection. Inflammation focal (red arrow). Magnification 100x.

## Discussion

COVID-19, an acute contagious respiratory disease caused by SARS-CoV-2 with high infectivity, is a worldwide concern. Clinical course of the disease showed that patients experienced a latent infection for one to two weeks, with high possibility to transmit the infection to other individuals (8). This suggests a possible distinct infection process of this virus. Most respiratory viruses, like influenza and rhinovirus, *in vitro* elicit a typical CPE in virus-infected cells (9,10), while SARS-CoV-2 does not (11). The lateral study noticed continuous release of virus particles from the human bronchial epithelial cells without cell damage. We have made similar observations in HSAEC used as a model epithelial cell line in our study. Collectively, these data show that SARS-CoV-2 uses a “clever” infection strategy that might be linked to the asymptomatic disease. This means that suitable preventive countermeasures should be considered. Therefore, antiviral drugs slowing down the replication of SARS-CoV-2 would be clinically beneficial.

Larifan as an antiviral drug, at the time of the invention, has been shown effective against many different viruses. Larifan inhibited the reproduction of cytomegalovirus (12) and replication of human immunodeficiency virus-1 (13) in cell cultures. The drug worked in immunocompromised mice infected with herpes simplex virus (HSV)-1 and HSV-2 (14) and prolonged the life span of monkeys infected with small pox (15). The drug demonstrated high antiviral efficacy against Omsk hemorrhagic fever in laboratory animals (16) and was effective against influenza (17) and rabies (18). In clinical setting, Larifan appeared effective against acute herpetic stomatitis in children (19), genital papillomavirus infections in women (20) and combined with herpetic vaccine led to improvement of the clinical symptoms of recurrence (21). Moreover, since its launch for use in human patients, it has been successfully used in the clinic predominantly against HSV infections with a safety track record.

Here, we demonstrate that Larifan reduces the replication of SARS-CoV-2 both *in vitro* and *in vivo* in SARS-CoV-2 infection model of Golden Syrian hamsters. Although we have observed an inhibitory effect of Larifan on SARS-CoV-2 replication, the underlying mechanism of antiviral effects is still elusive. We propose that Larifan mimics viral effects targeting the host and preventing further virus replication, including invading viruses and, thus, might have a dual action. First, direct antiviral action realized thought activation of enzymes of IFN system that leads to the global translation blockage of both cellular and viral mRNA in virus infected cells and, second, thought interfering with the host immune reaction (22,23). For that reason, we have chosen two different ways of Larifan administration, namely, i.n. that represents local and more direct mode of action and s.c. that corresponds to systemic effect. It is worth pointing out that i.n. admission of Larifan inhibited the virus infection better than the systemic administration in the s.c experiment model. These results might suggest that Larifan acts more locally in case of SARS-CoV-2 infection and could be potentially administered in a non-invasive manner *e*.*g*. in the form of nasal spray treatment. In this form, the medication could be utilized for preventative measures in high-risk people, and also as a treatment to reduce the virus numbers in infected individuals, especially in the early stages of the COVID-19 as it would promote faster recovery.

In conclusion, the inhibition of SARS-CoV-2 replication in *vitro* and the potential reduction of the viral load in the lungs of hamsters treated with Larifan alongside with improved lung histopathology, suggests towards the potential benefit of this drug in humans. However, further clinical studies are needed to confirm its effectiveness and safety in humans suffering from the novel coronavirus disease.

## Acknowledgements

The study was funded by Latvian Council of Science as fundamental and applied research project No lzp-2020/2-0369.

